# Cortical bone gain following loading is site-specifically enhanced by prior and concurrent disuse in aged male mice

**DOI:** 10.1101/705640

**Authors:** Gabriel Galea, Peter J Delisser, Lee Meakin, Lance E Lanyon, Joanna S Price, Sara H Windahl

## Abstract

The primary aim of bone anabolic therapies is to strategically increase bone mass in skeletal regions likely to experience high strains. This is naturally achieved by mechanical loading of the young healthy skeleton. However, these bone anabolic responses fail with age. Here, we applied site specificity analysis to map regional differences in bone anabolic responses to axial loading of the tibia (tri-weekly, for two weeks) between young (19-week-old) and aged (19-month-old), male and female mice. Loading increased bone mass specifically in the proximal tibia in both sexes and ages. Young female mice gained more cortical bone than young males in specific regions of the tibia. However, these site-specific sex difference were lost with age such that bone gain following loading was not significantly different between old males and females. Having previously demonstrated that prior and concurrent disuse enhances bone gain following loading in old females, we established whether this “rescue” is sex-specific. Old male mice were subjected to sciatic neurectomy or sham surgery, and tri-weekly loading was initiated four days after surgery. Disuse augmented cortical bone gain in response to loading in old male mice, but only in the regions of the tibia which were load-responsive in the young. Increased understanding of how locally-activated load-responsive processes lead to site-specific bone formation, and how the age-related diminution of these processes can be site-specifically enhanced by disuse, may lead to the next generation of strategic bone anabolic therapies.

**Highlights:**

- Sex differences in cortical tissue area of young and old mice are not site-specific
- The loading response in young, but not old, mice is sex- and site-specific
- The cortical loading response is site-specifically enhanced by disuse in old mice of both sexes
- The trabecular loading response can be rescued by disuse in old male, but not female, mice

## 1. Introduction

Sex differences in bone mass and architecture are well established in mice, humans, and other species [1–8]. Male bones are generally wider than female bones, and trabecular BMD in male mice is higher in the vertebrae but lower in the tibia compared to female mice [2]. Bone loss during ageing is different between the genders [5, 7]. During ageing, both men and women loose cortical bone endosteally, but because men also add more bone periosteally, the total bone loss is higher in women than in men [7]. Similarly, the trabecular bone loss during ageing is sex-specific. While the trabeculae in men become thinner with ageing, whole trabeculae are lost in women [5, 7].

Bone adapts to increased habitual loading by mainly increasing bone formation, a process that improves bone mass and architecture, and ultimately increases bone strength proportional to the load. These homeostatic processes are commonly referred to as the mechanostat [9–11]. When the mechanostat is compromised (e.g., in the elderly), fragility fractures increase [11]. When the tibiae of young adult mice are subjected to short periods of artificial loading, new bone is mainly deposited on the periosteal surface; however, this increase is greatly diminished in old mice [12–14].

Not only does male and female bone have a different appearance, they also respond differently to load [15–23]. Because female mice do not fight, while male mice fight every hour when housed in groups, female mice are more susceptible to increased loading than male mice [24]. In addition, male and female mice respond to load by partly using different mechanisms, where for example estrogen receptor α (ERα) is essential for the mechanostat in adult female but not male mice [21, 25].

Similarly, the mechanisms for the compromised mechanostat in old mice are sex-specific. Although the increased proliferation in response to load is blunted in both males and females, the cause for this effect is different. Osteoblasts from old male mice do not enter the cell-cycle in response to load, while female osteoblasts enter the cell-cycle but are arrested in the G2-pahse [12, 14]. Previously, we have shown that prior and concurrent disuse induced by sciatic neurectomy rejuvenates cortical bone of old female mice; this rejuvenation restores the loading response to levels found in young females [26]. Whether disuse similarly augments the response to loading in old male mice is not known.

These previous studies, investigated how bone responds to loading at only a few sites. To address this limitation, we have analyzed μCT images from the entire length of tibiae using site-specificity analysis (SSA) [27]. Using SSA, we have previously shown in young female mice that in response to load, cortical area increases at specific sites along the tibia [27]. In contrast, cortical bone decreases during ageing in female mice. Here, we applied SSA to investigate how sex and age influence bone mass, and how bone responds to loading, along the whole length of tibiae. In addition, we used SSA to determine whether disuse improves the bone response to loading in old male as it does in female mice.

## 2. Material and Methods

### 2.1 Animals

Young (19-week-old) and old (19-month-old) female and male C57BL/6J mice were obtained from Charles River Inc (Margate, UK) and housed in a standard animal facility under controlled temperature (22°C) and photoperiod (12 h of light, 12 h of dark) and fed pellet diet containing 0.75% calcium (EURodent Diet 22%; PMI Nutrition International, LLC, Brentwood, MO, USA) *ad libitum*. All procedures complied with the UK Animals (Scientific Procedures) Act 1986 under a UK Government Home Office project license (PPL30/2829) and were reviewed and approved by the University of Bristol ethics committee (Bristol, UK).

### 2.2 Ex vivo strain measurements

The magnitude of axial mechanical strain applied to the tibia during loading was established *ex vivo* to facilitate application of similar magnitudes of peak strain to all groups of mice. Sciatic neurectomy (SN, N=6) or sham (N=6) surgery was performed on the right limb as previously described [28, 29]. The mice were killed four days post-surgery and directly used for the *ex vivo* strain measurements as previously described [30, 31]. Strains were measured across a range of peak compressive loads between 4 and 18N (Supplemental Fig. 1). These peak loads were applied with the same ramped trapezoidal waveform, using the same 3100 ElectroForce® Test Instrument (Bose Corporation, MN, USA) with the same holding cups that were used later for *in vivo* loading. From the data, a linear regression analysis was performed. The strain:load relationship was not significantly altered four days after the sham or SN surgeries (Supplemental Fig. 1A). All old mice were therefore loaded with the same peak load during the experiment (14.5 N) to apply 2270με at the start of the loading experiment.

**Supplemental Figure S1.**
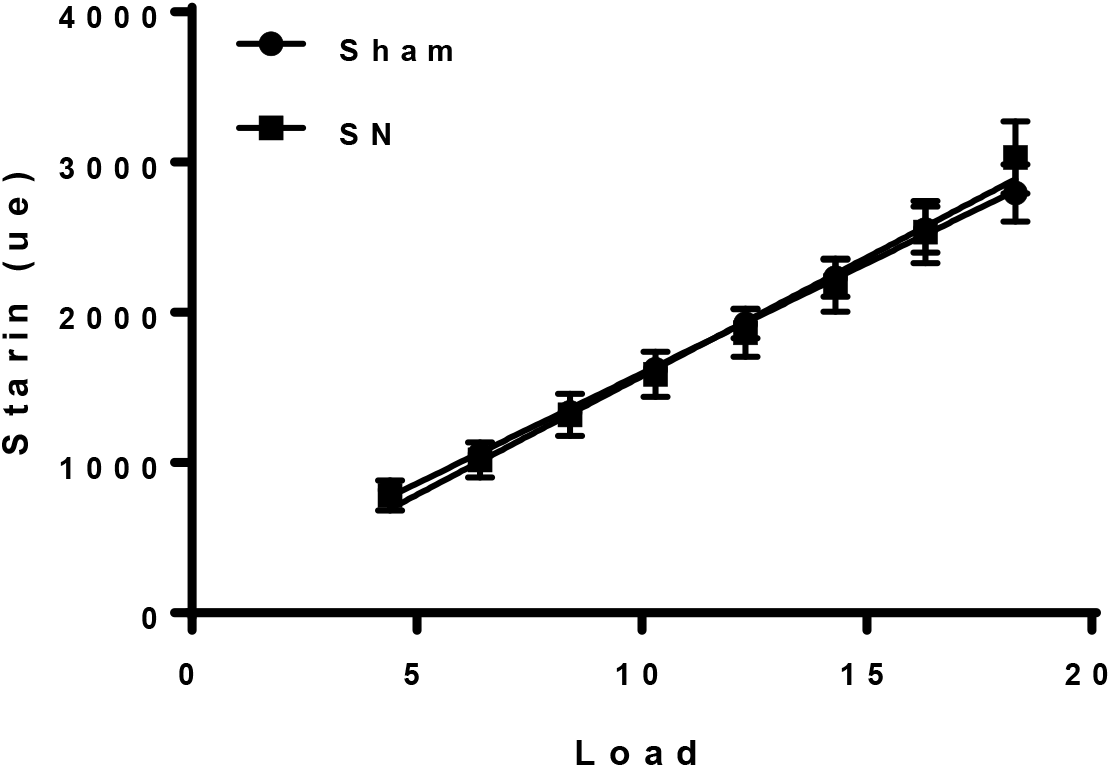
Four days unloading does not affect bone strength. Strain:load curves of tibiae 4 days post sham or neurectomy (SN) surgery of old male mice.

### 2.3 Mechanical loading

The protocol for non-invasively loading the mouse tibia has been reported previously [30–33]. Young male and female C57BL/6J mice (n = 6 per group) were subjected to loading so as to engender a peak strain of 2,500 με on the medial tibial surface [14]. In short, right tibias were subjected to external mechanical loading under isoflurane-induced anesthesia on alternate days for 2 weeks and mice were sacrificed 3 days following the last period of loading. All mice were allowed normal cage activity between loading sessions.

To determine whether disuse alters the response to loading in old males as is does in females [26], 19-month-old C57BL/6J male mice were divided in two weight-matched groups. The first group (N=13) was subjected to bilateral sham surgery and unilateral loading of the right tibia to demonstrate the effect of loading on a background of normal locomotion. The second group (N=12) was subjected to unilateral sham (left leg) and unilateral SN (right leg) surgery followed by loading of the right tibia to demonstrate the effect of loading on a background of disuse. All mice were kept in separate cages from the day of surgery so that fighting could not affect the loading response [24]. Four days post-surgery, the right tibias were subjected to external axial mechanical loading, every second day (in total 8 periods of loading), under isoflurane-induced anesthesia, to induce bone formation. Left limbs were used as internal controls as previously validated [32, 33]. An SN-only control group was deemed unnecessary given SN is well established to cause loss of bone mass in the disused limb [14, 26, 29, 34]. Effective SN was demonstrated by loss of muscle mass in each mouse. The mice were killed on the third day after the last loading, and the tibiae were dissected and fixed in 4% PFA for 48 hours. The tibiae were then stored in 70% EtOH until analysis by micro computed tomography (μCT). Female 19-month-old C57BL/6J mice were subjected to the same SN and loading protocol as the male mice [26].

### 2.4 High resolution μCT measurements of standard bone measurements

Whole tibiae were imaged using the SkyScan 1172 (Bruker, Kontich, Belgium) with a voxel size of 4.8μm (110μm^3^). The scanning, reconstruction and method of analysis has been previously reported [24, 35].

### 2.5 Site Specificity Analysis

SSA was performed and analyzed as previously described [27] and statistically analyzed using linear mixed models in SPSS (IBM, v.22). Bone site was used as a fixed categorical parameter, the intervention (loading, surgery) as a fixed effect, and an intervention by site interaction to account for site-specific responses. Mouse ID was included as a random effect with repeated measures at different sites from each mouse accounted for in the model. When the effect of the intervention was significant overall, a post-hoc Bonferroni correction was applied to identify individual sites where differences reached statistical significance. P < 0.05 was considered significant, but when several contiguous sites each tended towards significance (p < 0.01), these are also indicated.

### 2.6 Statistical analysis

Data is presented as mean ± SEM or as individual data points. Except when using SSA (statistical analysis described above), to evaluate the effect of loading (right loaded vs. left non-loaded control), surgery (sham vs. SN) and their interaction “loading*surgery”, a repeated-measures ANOVA was used with a post-hoc, paired t-test within groups. The effects of body weight, quadriceps wet weight and % changes to loading were also evaluated by paired t-tests (left versus right within each mouse). An effect was considered significant at p < 0.05.

## 3. Results

### 3.1 The sex differences in cortical tissue area of young and old mice are not site-specific

We first compared cortical bone of young and old, male and female mice using SSA. In both young and old mice, total tissue area (Tt.Ar) was greater in male than female mice in a non-site specific manner (sex x site interaction p = 0.991 and p=0.61, Fig. 1A and D respectively). Cortical area (Ct.Ar) was similarly significantly larger in male mice compared to female mice along most of the length of the tibia independent of age (Fig. 1B and E). However, the magnitude of this difference was site specific in the young (interaction p = 0.023), but not old (p = 0.52) mice.

**Figure 1.**
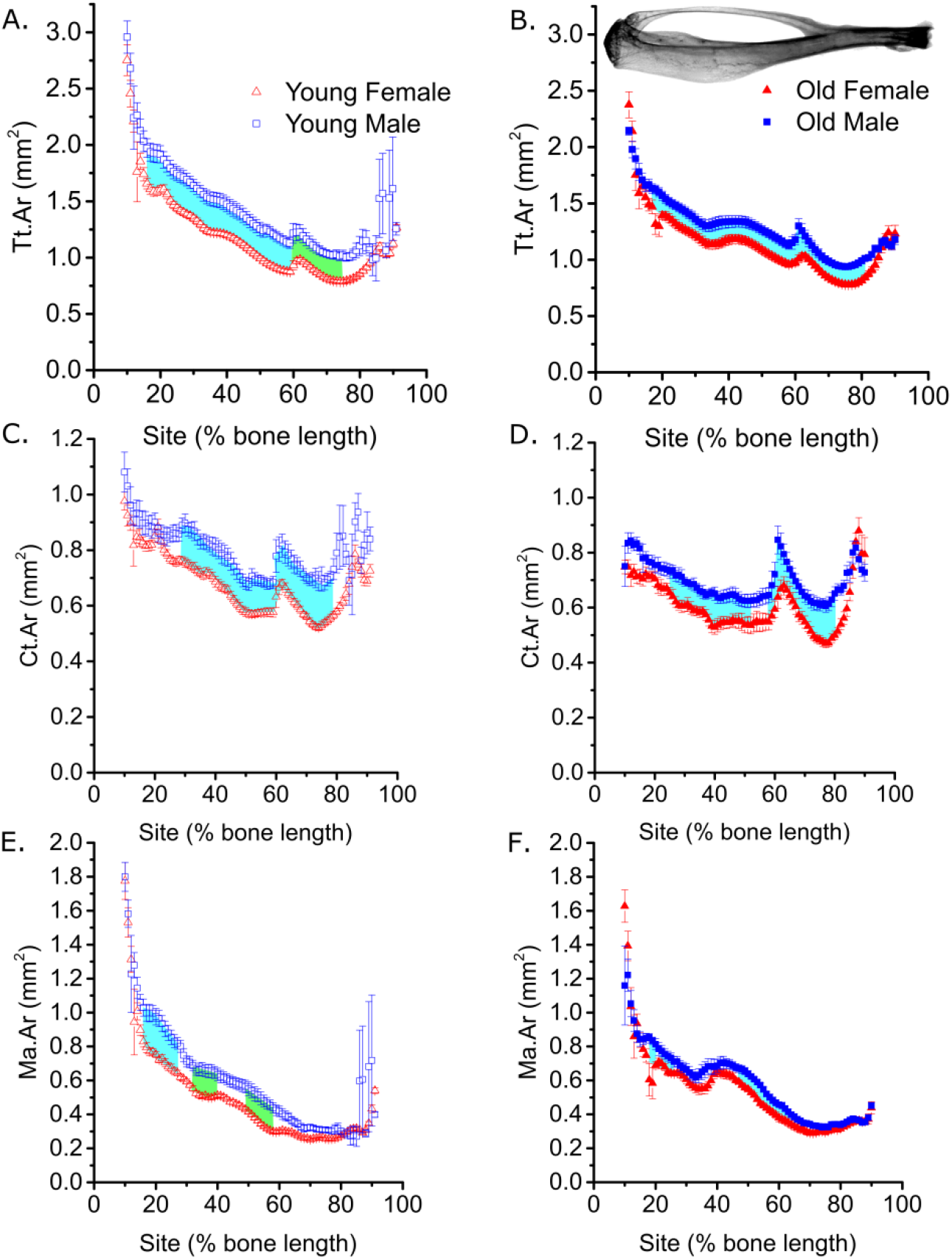
The sex differences in cortical tissue area of young and old mice are not site-specific. Site-specificity analysis of (A and b) periosteally enclosed area (Tt.Ar), (C and D) cortical area (Ct.Ar), (F and F) marrow area (Ma.Ar), of young (A, C and E) and old (B, F and F) male and female. Points represent the mean ± SEM, shaded regions indicate regions of statistical difference. N=6-7 and N=12-13 for young and old mice respectively. Turquoise shading p<0.05, green shading p<0.01.

Also independent of age, the marrow area (Ma.Ar) was larger in tibiae from males compared with females primarily in the proximal diaphysis, but not detectably so in the distal tibia, where the marrow cavity is small (Fig. 1C and F). The difference in Ma.Ar was not significantly site-specific in the young (interaction p = 0.470), but was significantly site-specific in the old (p = 0.04).

### 3.2 The loading response in young, but not old, mice is sex- and site-specific

In young male and female mice, Tt.Ar increased following loading primarily in the proximal tibia (site * loading interaction p < 0.001 for both sexes). There was no significant difference in the loading response in Tt.Ar in male compared to female mice (overall p = 0.75 comparing male vs. female), although female mice tended to show a greater percentage increase on average around 37% of the bone’s length from the proximal end (Fig. 2A). The loading-related increase in Ct.Ar was also site-specific in both sexes (site * loading interaction p < 0.01), and was significantly greater in the proximal to central diaphysis of the female compared to the male tibiae (Fig. 2B). In females, a site of maximum responsiveness was observed around the 37% site, as we have previous described [27]. However, in young male mice, no such “peak” was observed (Fig. 2B). Loading was also associated with a greater reduction in Ma.Ar in female than male mice at 30-36% and 60-68% of the bone’s length from the proximal end (Fig. 2C). Taken together, these findings demonstrate that the adaptive cortical responses following axial loading are site-specific in both sexes, but are more marked and follow a different spatial pattern in female than male mice subjected to the same peak strain stimulus. In old mice, the loading-related change in Tt.Ar (Fig. 2D), Ct.Ar (Fig. 2E) and Ma.Ar (Fig. 2F) was not significantly different overall between males and females.

**Figure 2.**
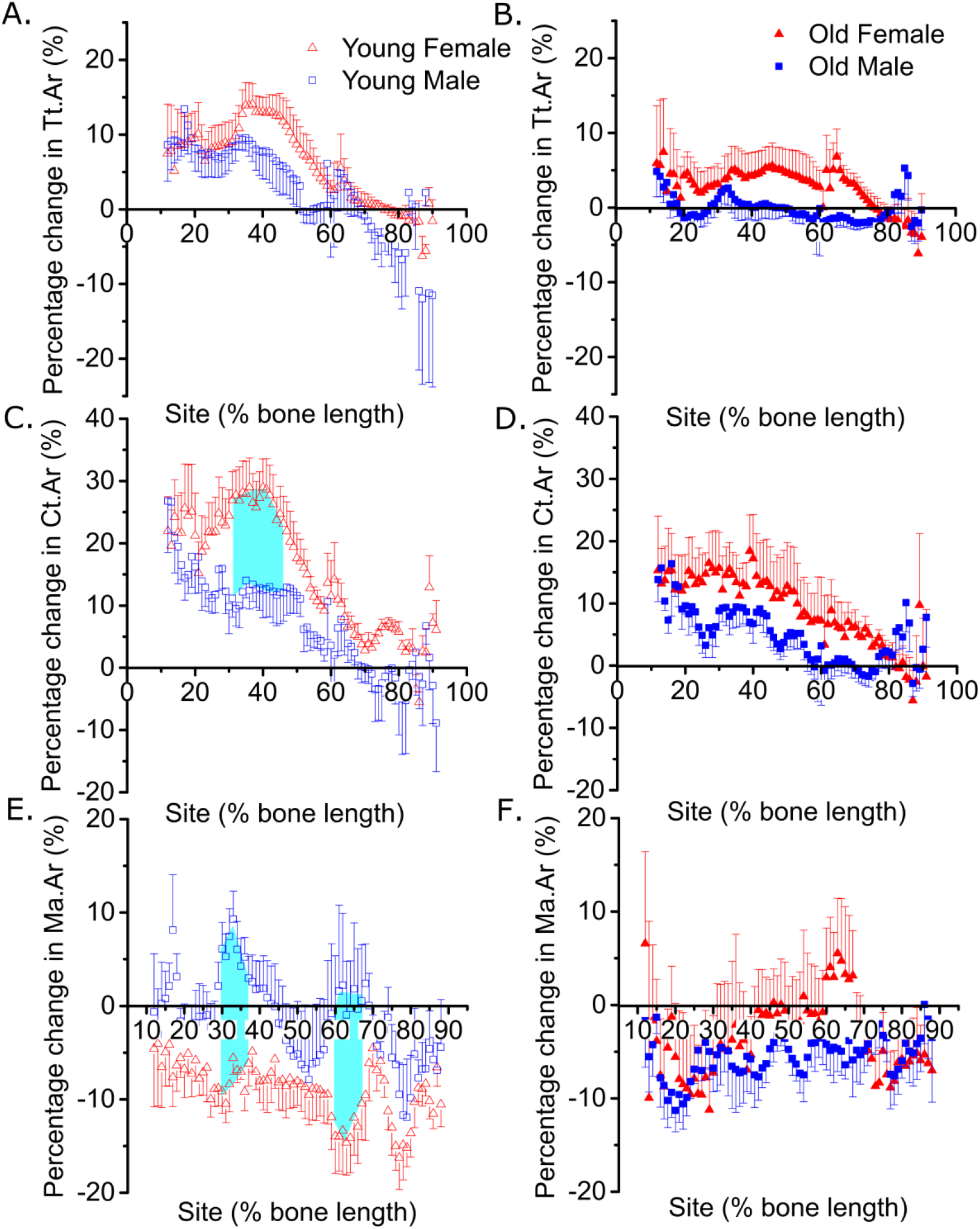
The loading response in young, but not old, mice is sex- and site-specific. Site-specificity analysis of the loading-related percent change in (A and B) periosteally enclosed area (Tt.Ar), (C and D) cortical area (Ct.Ar), (E and F) marrow area (Ma.Ar) of young male and female (A, C and E), and old male and female (B, D and F) mice. Points represent the mean ± SEM, shaded regions indicate regions of statistical difference. N=6-7 and N=12-13 for young and old mice respectively.

### 3.3 The decreased loading response in old mice is sex- and site-specific

In old mice, the overall loading-related increase in Tt.Ar and Ct.Ar was confirmed to be smaller than in young mice of both sexes as we have previously reported [14]. However, SSA provided substantial additional information. The loading-related increase in Tt.Ar and Ct.Ar tended to be diminished in the old versus young females specifically around the peak responsiveness site, approximately 37% of the bone’s length from the proximal end (Fig. 3A and B). However, in males, a significant age-related difference in the loading-related change in Tt.Ar and Ct.Ar was detected more proximally, at around 20% of the bone’s length (Fig. 3D and E) and not the commonly-analyzed 37% or 50% sites. Ma.Ar decreased following loading in both young and old-female mice in the proximal tibia, but the reduction was more marked distally in the young mice as we have previously reported [27]. This distal reduction in Ma.Ar was not observed in old female mice (Fig. 3C). Old male mice showed significantly greater reductions in Ma.Ar than young mice and this difference was again most marked in the proximal tibia around the 20% site (Fig. 3F).

**Figure 3.**
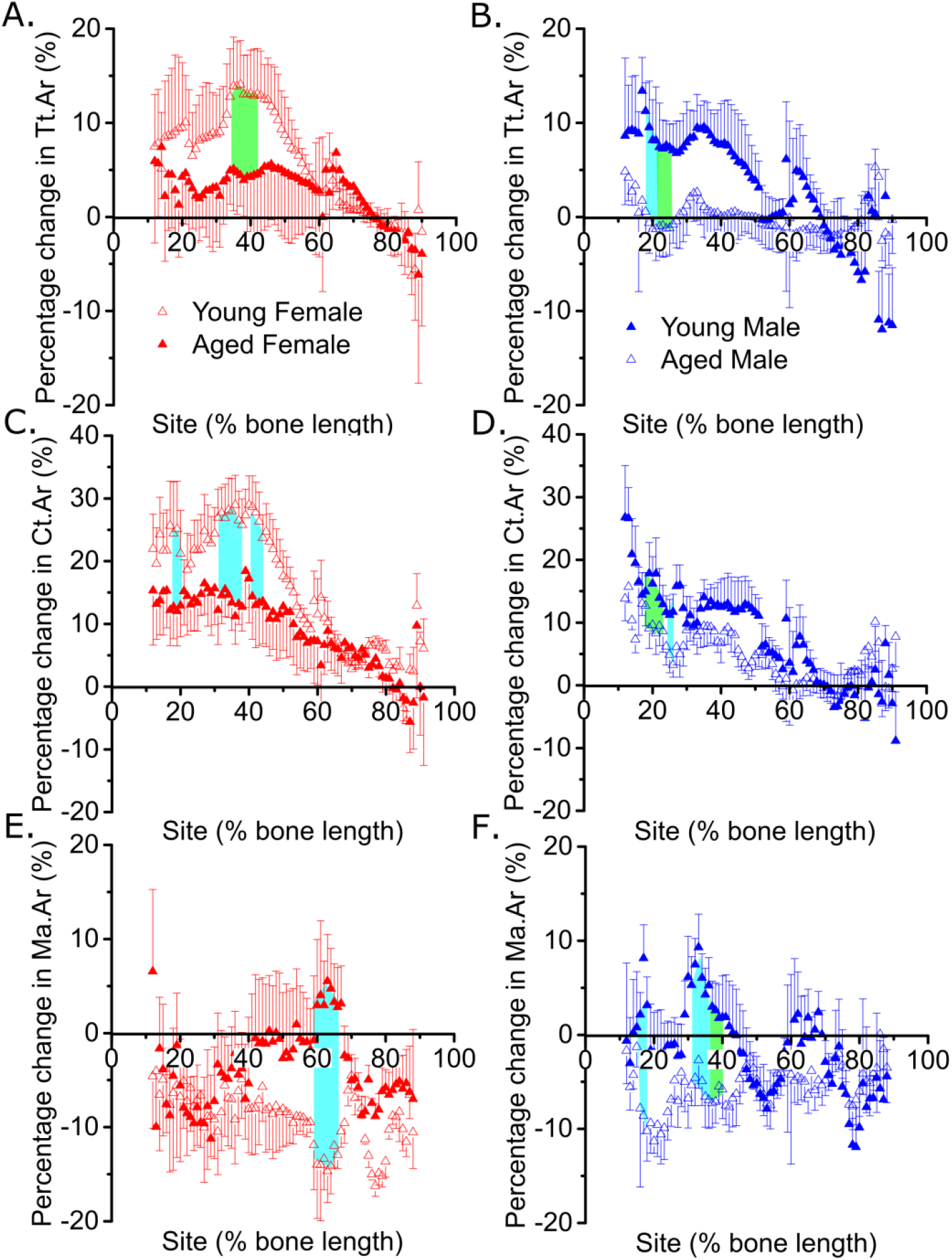
The decreased loading response in old mice is sex- and site-specific. Site-specificity analysis of the loading-related percent change in (A and B) periosteally enclosed area (Tt.Ar), (C and D) cortical area (Ct.Ar), (E and F) marrow area (Ma.Ar) of young and old female (A, C and E) and young and old male (B, D and F) mice. Points represent the mean ± SEM, shaded regions indicate regions of statistical difference. N=6-7 and N=12-13 for young and old mice respectively

### 3.4 Disuse site-specifically enhances the cortical adaptive responses to loading in old males

We have previously shown that prior and concurrent disuse can “rescue” the age-related decline in cortical bones’ response to loading in female mice. To establish if this response is sex-specific, we used SSA to investigate the effect of prior and concurrent disuse on the loading response in old male mice. SN significantly reduced quadriceps wet weight in the right leg (−41%, p<0.001) compared to the left control leg of SN mice (Supplemental Figure 2A), confirming that the SN was successful. The overall body weight was not affected by SN (Supplementary Figure 2B).

**Supplemental figure 2.**
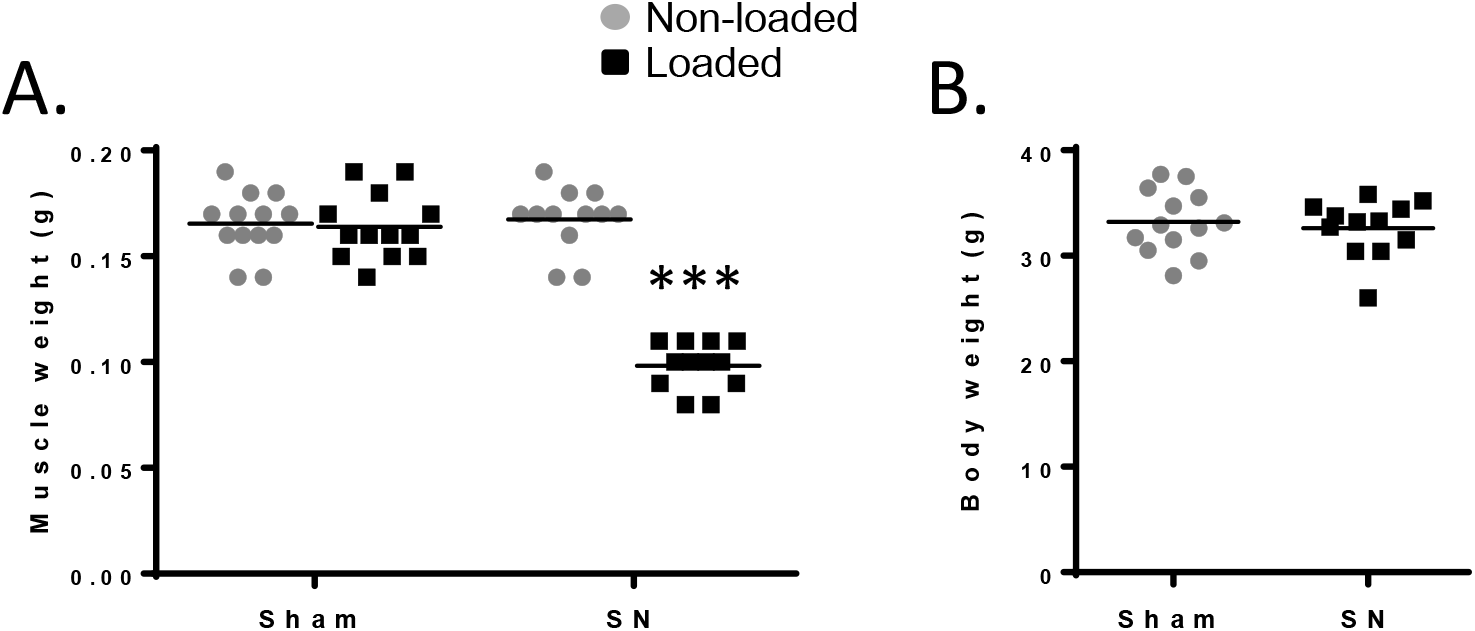
Muscle weight, but not body weight is affected by neurectomy. (A) Gastrocnemius and (B) body weights are shown as mean ± SEM for sham and neurectomized (SN) mice. ***p<0.001 for non-loaded vs. loaded tibiae of the same treatment.

Loading on a background of disuse resulted in a significantly greater increase in Tt.Ar predominantly around the 20% site in old males (Fig. 4A and 6A), the region which showed the greatest age-related decline in responsiveness. Similarly, the increase in Ct.Ar was significantly greater in SN-than sham-operated mice in the proximal tibia, including significant enhancement at the 20% site (Fig. 4B and 6B). In contrast, the reduction in Ma.Ar observed at ~20-30% of the bone’s length in old male mice was not significantly altered by disuse (Fig. 4C).

**Figure 4.**
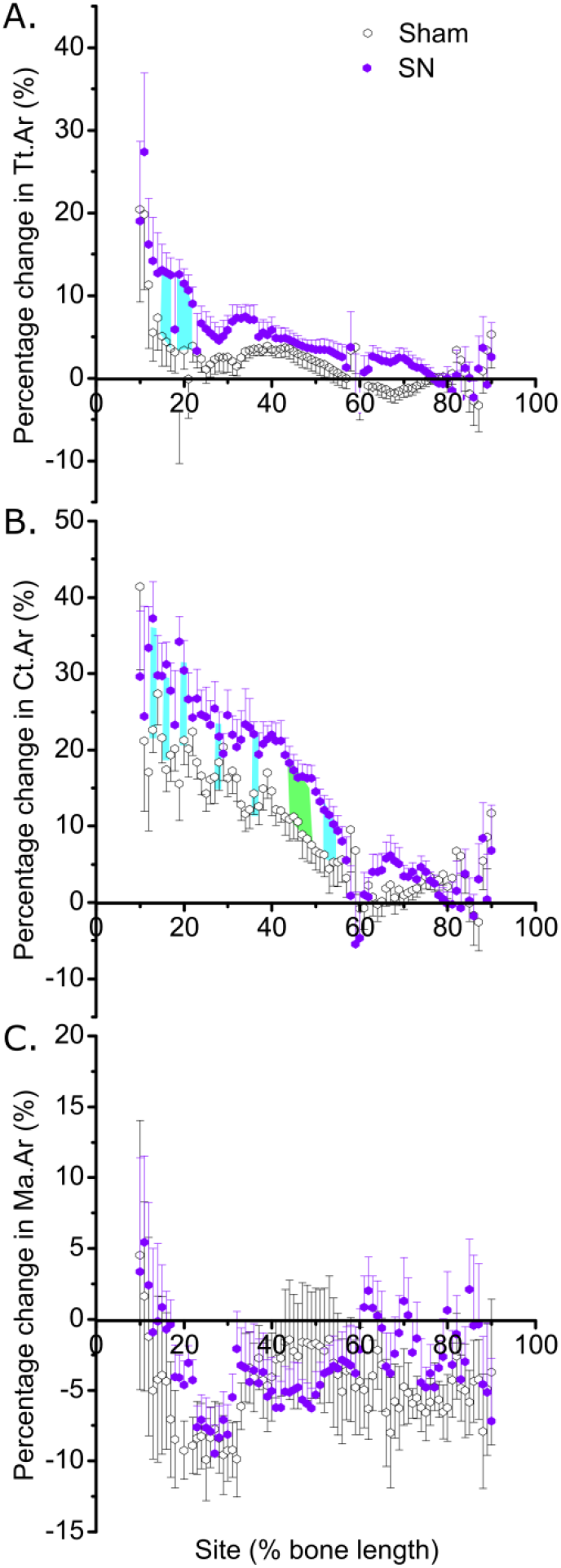
Concurrent disuse rescues the age-dependent reduced loading response in male mice. Site-specificity analysis of the loading-related percent change in (A) periosteally enclosed area (Tt.Ar), (B) cortical area (Ct.Ar), (C) marrow area (Ma.Ar) of sham and SN old male mice. Points represent the mean ± SEM, shaded regions indicate regions of statistical difference. N=13 for sham and N=12 for SN. Turquoise shading p<0.05, green shading p<0.01.

These findings were validated by conventional μCT analysis selectively at the cortical 20% site (Figures 5A and 6, and Table 1). Other standard measures of cortical bone mass were also assessed. Cortical cross-sectional thickness (Cs.Th, Table 1) and cortical area/periosteally enclosed area (Ct.Ar/Tt.Ar, Table 1) increased following loading irrespective of SN, whereas polar moment of inertia (PMI, Figure 6C) increased to a greater extent in the SN versus Sham group relative to the left limbs. Thus, prior and concurrent disuse enhances the cortical osteogenic response to loading in old male mice as we have previously reported in old females [26]. This “rescue” is site-specific and most pronounced at the cortical 20% site in males, where it was readily validated by conventional μCT analysis (Figures 5A and 6, and Table 1).

**Figure 5.**
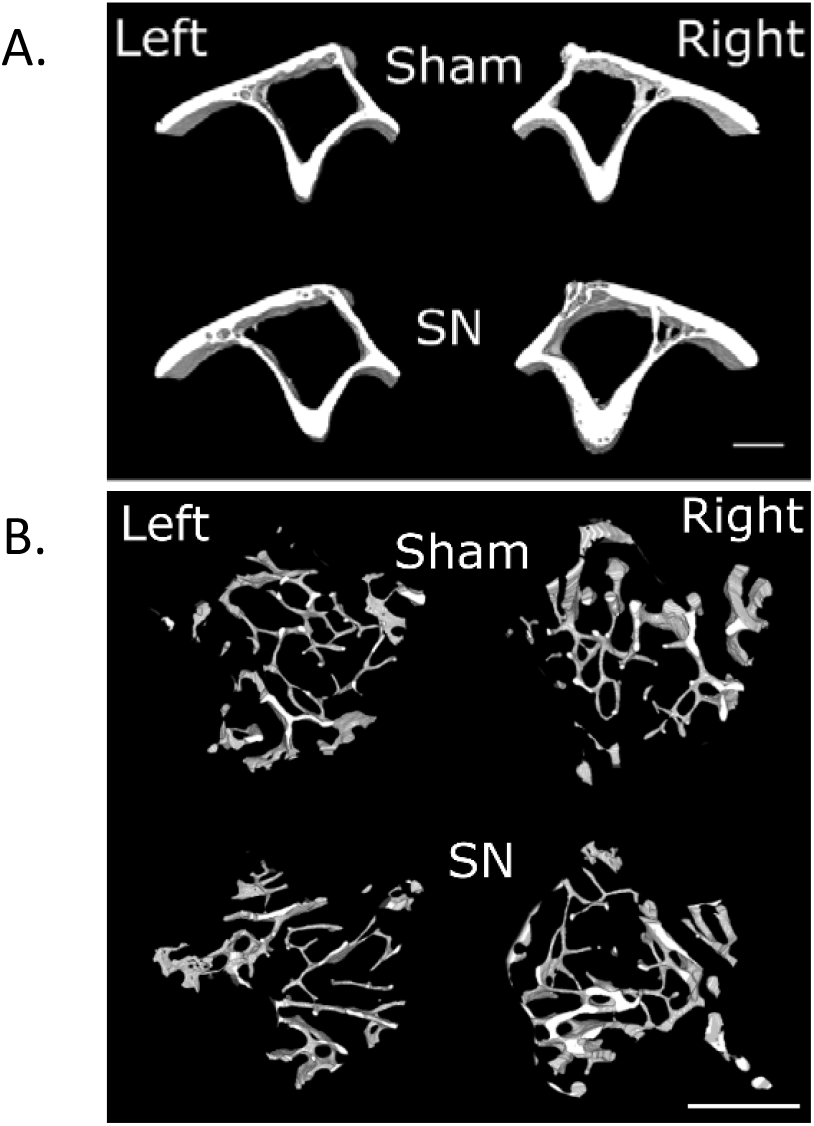
Representative μCT images. Right legs were subjected to axial loading, while the left legs are internal controls. SN = sciatic neurectomy.

**Figure 6.**
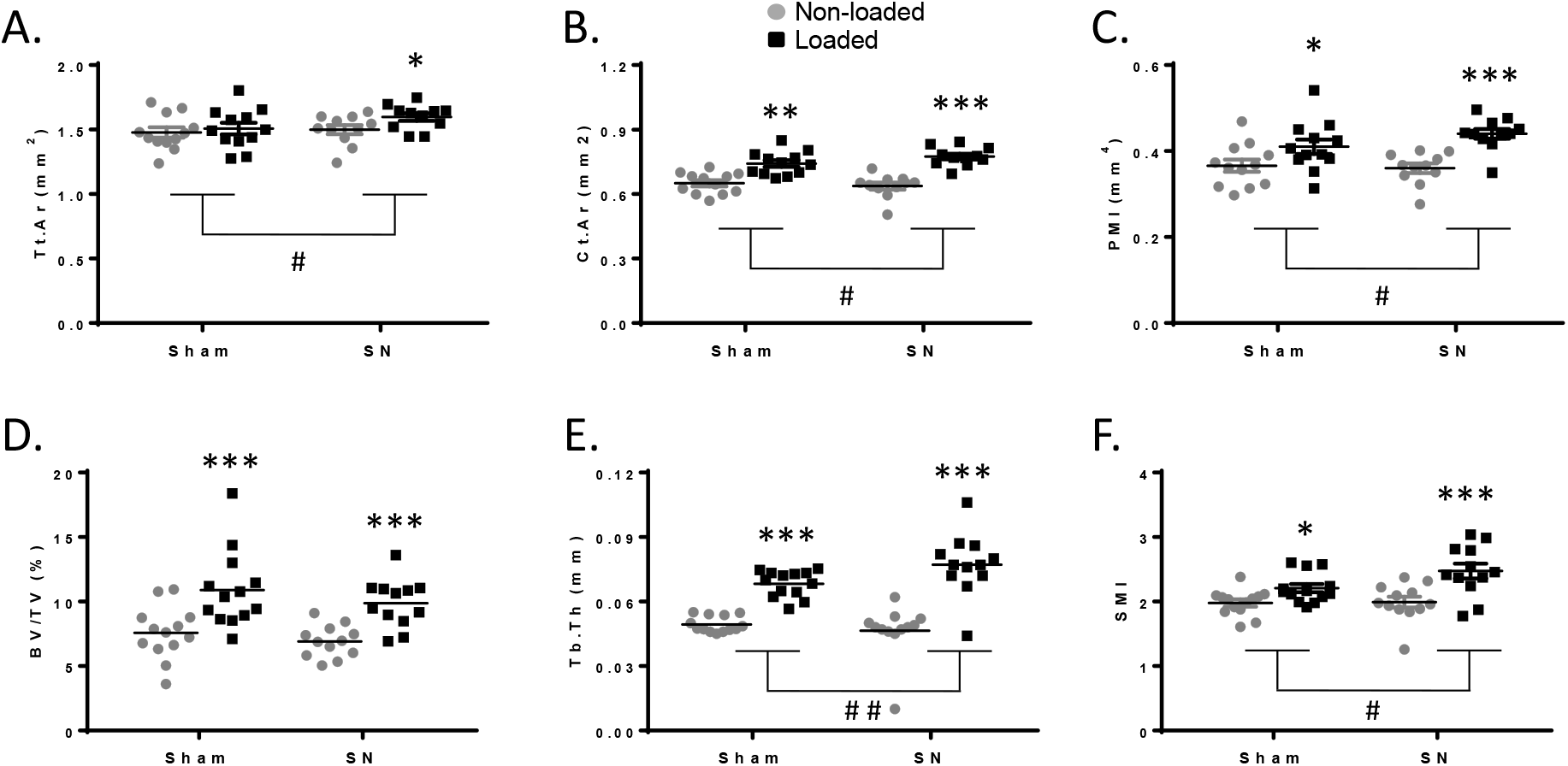
Conventional μCT confirms the SSA analysis. Conventional μCT analysis of (A) periosteally enclosed (Tt.Ar), (B) cortical area (Ct.Ar), (C) polar moment of inertia (PMI), (D) trabecular bone volume (BV/TV), (E) trabecular thickness (Tb.Th) and (F) structure model index (SMI) are shown as mean ± sem for sham and neurectomized (SN) mice. Left non-loaded tibiae are represented by white boxes and right loaded tibiae are indicated by black boxes. N=13 for sham and N=12 for SN.

### 3.5 Disuse enhances the trabecular adaptive responses to loading in old male mice

We also investigated the effect of prior and concurrent disuse on the load-induced effects on trabecular bone. Loading significantly increased trabecular bone volume fraction (BV/TV), trabecular thickness (Tb.Th) and structure model index (SMI, Figure 5B and 6D-F), but decreased the trabecular patterning factor (Tb.Pf, Table 1) in both sham and SN tibiae of old male mice. Trabecular number (Tb.N) and trabecular separation (Tb.Sp) did not change in either group (Table 1). Prior and concurrent disuse significantly augmented the loading-induced increase in Tb.Th and PMI (Figure 6E and F).

## 4. Discussion

Male bones are generally larger than female bones, and the shape of some bones such as the pelvis are also sex-specific. Irrespective of sex, non-uniform structural and material adaptations to loading influence bone architecture. Yet, previous studies on bone adaptation to loading have only investigated a few sites. To address this limitation, we used SSA to analyze μCT images along the entire length of mouse tibiae. As expected, the diameter of the tibiae is larger along most of the tibiae in both young and old males compared to females. This difference in diameter is attributed to several factors including diet, hormones, body composition, and mechanical loading (e.g., male mice fight more often) [5, 15, 24]. We show that, despite sex differences in diameter size, there are similarities in tibial architecture between the sexes; for example, the tibia/fibula junction (shown by the inflection in Ct.Ar) is located at the same site in young and old mice of both sexes.

We found that in young mice the loading response differed in magnitude and distribution between the sexes. These findings are consistent with previous studies showing a larger load-related increase in cortical bone mass in young female mice compared to young male mice at the 37% site [14, 21]. However, it is not simply the case that male mice are generally less responsive than female mice as loading-related changes in cortical parameters were comparable between young males and females at the proximal end of the tibiae. In other words, we found that young female mice had a larger load-related increase in cortical bone mass at specific sites. Because males and females were subjected to the same peak strain at the 37% site, the differences in responsiveness suggest that the site-specificity of the loading response is not only affected by the strain induced, but also by other factors. These factors may include estrogen and androgen receptors, which affect the loading response differently in the two sexes [15, 21]. Although it has previously been demonstrated that the osteogenic responses to loading at individual cortical sites could be affected by genetic manipulations or pharmacological treatments (e.g., androgens, intermittent parathyroid hormone, and selective estrogen receptor modulators) [36–39], it remains to be determined whether these interventions also result in local changes of the loading response.

We have previously shown that the loading response is site-specific in young and old female mice [27]. Here, we show that the loading response is also site-specific in young and old male mice. The loading response is stronger proximally in young males compared to old males, whereas the loading response is similar more distally in young males and old males. Ageing diminishes the loading-induced increase in cortical area mainly due to diminished periosteal expansion in the proximal tibiae in both male and female mice [14]. This study shows that sexual differences in the cortical loading response are less pronounced in old male and female mice. In addition, none of the loading-related changes in any of the standard cortical parameters measured were significantly different between old male and female mice. Because the loading responses in old mice were more variable, we cannot exclude the possibility that undetectable differences in loading response may exist between the sexes. However, the spatial distributions of responses observed also followed a similar pattern in old males and females. This regression towards similar responsiveness to loading in old mice could be related to one or several changes related to ageing. One of these changes could be cellular senescence, which increases with age. This increase is associated with decreased bone mass and bone formation, and when senescent cells are removed or reversed, bone mass is increased in old mice [40]. Further research is needed to determine which mechanisms govern the regression towards similar responsiveness to loading seen in old mice.

The magnitude of site-specific responses to mechanical loading is influenced by the systemic (e.g., hormonal and genetic) osteogenic context of the loading. We have recently proposed that this osteogenic context is also locally influenced by concurrent habitual loading stimuli (e.g., recurrent fighting among young males reduces the loading response) [24]. Also, the age-related impairment of the loading response can be, at least partly, rejuvenated by prior and concurrent disuse in old female mice [26]. In setting out to investigate whether disuse similarly enhances the loading response in old male mice, we were faced with the problem that, as we previously observed, loading-related changes in cortical parameters are not significantly different between young and old male mice when analyzed at the single commonly assayed 37% site [14].

Here, we show that the loading response is blunted further periosteally in old compared to young male mice, and that prior and concurrent disuse rejuvenates the osteogenic response to loading in old male mice. Specifically, prior and concurrent disuse rejuvenated periosteal expansion resulting in a greater increase in Tt.Ar, thereby augmenting the gain in overall Ct.Ar in old males. The mechanisms behind this rejuvenation in either sex are unknown, partly because the impairment of the mechanostat with age remains incompletely understood. Possible mechanisms for the reduced loading response in old mice include a blunted down-regulation of Wnt antagonists following loading in osteocytes (*in vivo*) [41] and reduced osteoblast proliferation following strain (*in vitro*) [14]. Microarray comparison of tibiae from young and old female mice suggests that cellular processes related to bioenergetics and cell proliferation (both of which are differentially expressed between habitually loaded limbs of the two age groups) also show divergent responses to additional loading [42]. Furthermore, many of the genes differentially regulated by loading are also differentially expressed during disuse [43]. Thus, prior and concurrent disuse may “prime” loading-responsive cellular processes, which are basally diminished with age. However, transcriptomic studies describing age- and loading-associated changes have not been able to discriminate between cortical sites. In addition to differences between cortical sites, there are several well-established differences between the determinants of trabecular and cortical bone mass and their response to loading, including estrogen and androgen receptors. For example, in young mice, deletion of ERα decreases cortical BMD but increases trabecular bone in male and female mice [22]. In contrast, deletion of ERβ increases cortical but not trabecular bone mass in young female, but not male, mice [22, 44, 45]. Further, ERα is essential for the cortical but not the trabecular response to loading in female mice but may inhibit the loading response in both the cortical and trabecular compartments in young male mice [21, 31]. In contrast, deletion of ERβ increases the cortical, but not trabecular, response to loading in both male and female mice [21]. Further research is needed to reveal how these receptors influence the loading response in old mice.

The trabecular loading response is also site-specific, as loading down-regulates sclerostin and increases bone volume in the secondary, but not primary spongiosa [28]. In old female mice, disuse did not alter the response of trabecular bone to loading [26], although the extremely low level of baseline trabecular bone may have prevented detection of any loading-related changes. Here, we show that disuse enhanced the load-related increase in Tb.Th and SMI in old male mice. Thus, the loading response of both cortical and trabecular compartments is sensitive to habitual levels of loading, further suggesting that the mechanostat regulates bone mass by monitoring loading over time as a way to adapt to changing environmental conditions. Age-related desensitization may account for the age-related impairment of the mechanostat.

Here, we applied SSA to map the skeletal sexual differences in cortical bone structure and loading responsiveness in young and old mice. Despite having a lower baseline bone mass, young female mice show osteogenic responses greater than young male mice in the most load-responsive region of the tibial diaphysis. However, these loading-response differences decrease with age such that the loading response of old female mice is comparable to that of old male mice. Prior and concurrent disuse rejuvenated the mechanostat in old males, augmenting the osteogenic response to loading. We propose that sex, age, and site should be considered when performing and comparing analyses of mouse bones. Furthermore, these results suggests that brief bouts of intense physical activity may increase bone mass in sedentary elderly men and women.

## 5. Acknowledgements

This study was supported by a European Union’s Horizon 2020 research and innovation program under the Marie Skłodowska-Curie grant agreement No 657178 (SW), Wellcome Veterinary Intercalated Training Fellowships (to GLG 088560/Z/09/Z), a Wellcome Postdoctoral Clinical Research Training Fellowship (to GLG, 107474/Z/15/Z), the Swedish Research Council (grant agreement number 2013-8252, SW).

## References

[1] T.J. Beck, C.B. Ruff, W.W. Scott, Jr., C.C. Plato, J.D. Tobin, C.A. Quan, Sex differences in geometry of the femoral neck with aging: a structural analysis of bone mineral data, Calcif Tissue Int 50(1) (1992) 24–9.

[2] E.A. Carson, J.P. Kenney-Hunt, M. Pavlicev, K.A. Bouckaert, A.J. Chinn, M.J. Silva, J.M. Cheverud, Weak genetic relationship between trabecular bone morphology and obesity in mice, Bone 51(1) (2012) 46–53.

[3] S. Kaptoge, N. Dalzell, N. Loveridge, T.J. Beck, K.T. Khaw, J. Reeve, Effects of gender, anthropometric variables, and aging on the evolution of hip strength in men and women aged over 65, Bone 32(5) (2003) 561–70.

[4] M. Laurent, L. Antonio, M. Sinnesael, V. Dubois, E. Gielen, F. Classens, D. Vanderschueren, Androgens and estrogens in skeletal sexual dimorphism, Asian J Androl 16(2) (2014) 213–22.

[5] J.W. Nieves, Sex-Differences in Skeletal Growth and Aging, Curr Osteoporos Rep 15(2) (2017) 70–75.

[6] A. Riesenfeld, Sexual dimorphism of skeletal robusticity in several mammalian orders, Acta Anat (Basel) 102(4) (1978) 392–8.

[7] E. Seeman, Clinical review 137: Sexual dimorphism in skeletal size, density, and strength, J Clin Endocrinol Metab 86(10) (2001) 4576–84.

[8] M. Sinnesael, F. Claessens, S. Boonen, D. Vanderschueren, Novel insights in the regulation and mechanism of androgen action on bone, Curr Opin Endocrinol Diabetes Obes 20(3) (2013) 240–4.

[9] H.M. Frost, The mechanostat: a proposed pathogenic mechanism of osteoporoses and the bone mass effects of mechanical and nonmechanical agents, Bone Miner 2(2) (1987) 73–85.

[10] H.M. Frost, Perspectives: a proposed general model of the “mechanostat” (suggestions from a new skeletal-biologic paradigm), Anat Rec 244(2) (1996) 139–47.

[11] L. Lanyon, T. Skerry, Postmenopausal osteoporosis as a failure of bone’s adaptation to functional loading: a hypothesis, J Bone Miner Res 16(11) (2001) 1937–47.

[12] M.D. Brodt, M.J. Silva, Aged mice have enhanced endocortical response and normal periosteal response compared with young-adult mice following 1 week of axial tibial compression, J Bone Miner Res 25(9) (2010) 2006–15.

[13] N. Holguin, M.D. Brodt, M.E. Sanchez, M.J. Silva, Aging diminishes lamellar and woven bone formation induced by tibial compression in adult C57BL/6, Bone 65 (2014) 83–91.

[14] L.B. Meakin, G.L. Galea, T. Sugiyama, L.E. Lanyon, J.S. Price, Age-related impairment of bones’ adaptive response to loading in mice is associated with sex-related deficiencies in osteoblasts but no change in osteocytes, J Bone Miner Res 29(8) (2014) 1859–71.

[15] F. Callewaert, M. Sinnesael, E. Gielen, S. Boonen, D. Vanderschueren, Skeletal sexual dimorphism: relative contribution of sex steroids, GH-IGF1, and mechanical loading, J Endocrinol 207(2) (2010) 127–34.

[16] G. Cardadeiro, F. Baptista, R. Ornelas, K.F. Janz, L.B. Sardinha, Sex specific association of physical activity on proximal femur BMD in 9 to 10 year-old children, PLoS One 7(11) (2012) e50657.

[17] R.M. Daly, The effect of exercise on bone mass and structural geometry during growth, Med Sport Sci 51 (2007) 33–49.

[18] S. Kriemler, L. Zahner, J.J. Puder, C. Braun-Fahrlander, C. Schindler, N.J. Farpour-Lambert, M. Kranzlin, R. Rizzoli, Weight-bearing bones are more sensitive to physical exercise in boys than in girls during pre- and early puberty: a cross-sectional study, Osteoporos Int 19(12) (2008) 1749–58.

[19] K.M. Melville, N.H. Kelly, G. Surita, D.B. Buchalter, J.C. Schimenti, R.P. Main, F.P. Ross, M.C. van der Meulen, Effects of Deletion of ERalpha in Osteoblast-Lineage Cells on Bone Mass and Adaptation to Mechanical Loading Differ in Female and Male Mice, J Bone Miner Res 30(8) (2015) 1468–80.

[20] L.D. Moreno, S.D. Waldman, M.D. Grynpas, Sex differences in long bone fatigue using a rat model, J Orthop Res 24(10) (2006) 1926–32.

[21] L.K. Saxon, G. Galea, L. Meakin, J. Price, L.E. Lanyon, Estrogen receptors alpha and beta have different gender-dependent effects on the adaptive responses to load bearing in cancellous and cortical bone, Endocrinology 153(5) (2012) 2254–66.

[22] N.A. Sims, S. Dupont, A. Krust, P. Clement-Lacroix, D. Minet, M. Resche-Rigon, M. Gaillard-Kelly, R. Baron, Deletion of estrogen receptors reveals a regulatory role for estrogen receptors-beta in bone remodeling in females but not in males, Bone 30(1) (2002) 18–25.

[23] S.H. Windahl, G. Andersson, J.A. Gustafsson, Elucidation of estrogen receptor function in bone with the use of mouse models, Trends Endocrinol Metab 13(5) (2002) 195–200.

[24] L.B. Meakin, T. Sugiyama, G.L. Galea, W.J. Browne, L.E. Lanyon, J.S. Price, Male mice housed in groups engage in frequent fighting and show a lower response to additional bone loading than females or individually housed males that do not fight, Bone 54(1) (2013) 113–7.

[25] F. Callewaert, A. Bakker, J. Schrooten, B. Van Meerbeek, G. Verhoeven, S. Boonen, D. Vanderschueren, Androgen receptor disruption increases the osteogenic response to mechanical loading in male mice, J Bone Miner Res 25(1) (2010) 124–31.

[26] L.B. Meakin, P.J. Delisser, G.L. Galea, L.E. Lanyon, J.S. Price, Disuse rescues the age-impaired adaptive response to external loading in mice, Osteoporos Int 26(11) (2015) 2703–8.

[27] G.L. Galea, S. Hannuna, L.B. Meakin, P.J. Delisser, L.E. Lanyon, J.S. Price, Quantification of Alterations in Cortical Bone Geometry Using Site Specificity Software in Mouse models of Aging and the Responses to Ovariectomy and Altered Loading, Front Endocrinol (Lausanne) 6 (2015) 52.

[28] A. Moustafa, T. Sugiyama, J. Prasad, G. Zaman, T.S. Gross, L.E. Lanyon, J.S. Price, Mechanical loading-related changes in osteocyte sclerostin expression in mice are more closely associated with the subsequent osteogenic response than the peak strains engendered, Osteoporos Int 23(4) (2012) 1225–34.

[29] T. Sugiyama, L.B. Meakin, W.J. Browne, G.L. Galea, J.S. Price, L.E. Lanyon, Bones’ adaptive response to mechanical loading is essentially linear between the low strains associated with disuse and the high strains associated with the lamellar/woven bone transition, J Bone Miner Res 27(8) (2012) 1784–93.

[30] H. Todd, G.L. Galea, L.B. Meakin, P.J. Delisser, L.E. Lanyon, S.H. Windahl, J.S. Price, Wnt16 Is Associated with Age-Related Bone Loss and Estrogen Withdrawal in Murine Bone, PLoS One 10(10) (2015) e0140260.

[31] S.H. Windahl, L. Saxon, A.E. Borjesson, M.K. Lagerquist, B. Frenkel, P. Henning, U.H. Lerner, G.L. Galea, L.B. Meakin, C. Engdahl, K. Sjogren, M.C. Antal, A. Krust, P. Chambon, L.E. Lanyon, J.S. Price, C. Ohlsson, Estrogen receptor-alpha is required for the osteogenic response to mechanical loading in a ligand-independent manner involving its activation function 1 but not 2, J Bone Miner Res 28(2) (2013) 291–301.

[32] R.L. De Souza, M. Matsuura, F. Eckstein, S.C. Rawlinson, L.E. Lanyon, A.A. Pitsillides, Non-invasive axial loading of mouse tibiae increases cortical bone formation and modifies trabecular organization: a new model to study cortical and cancellous compartments in a single loaded element, Bone 37(6) (2005) 810–8.

[33] T. Sugiyama, J.S. Price, L.E. Lanyon, Functional adaptation to mechanical loading in both cortical and cancellous bone is controlled locally and is confined to the loaded bones, Bone 46(2) (2010) 314–21.

[34] L.K. Saxon, B.F. Jackson, T. Sugiyama, L.E. Lanyon, J.S. Price, Analysis of multiple bone responses to graded strains above functional levels, and to disuse, in mice in vivo show that the human Lrp5 G171V High Bone Mass mutation increases the osteogenic response to loading but that lack of Lrp5 activity reduces it, Bone 49(2) (2011) 184–93.

[35] T. Sugiyama, L.B. Meakin, G.L. Galea, B.F. Jackson, L.E. Lanyon, F.H. Ebetino, R.G. Russell, J.S. Price, Risedronate does not reduce mechanical loading-related increases in cortical and trabecular bone mass in mice, Bone 49(1) (2011) 133–9.

[36] L.B. Meakin, H. Todd, P.J. Delisser, G.L. Galea, A. Moustafa, L.E. Lanyon, S.H. Windahl, J.S. Price, Parathyroid hormone’s enhancement of bones’ osteogenic response to loading is affected by ageing in a dose- and time-dependent manner, Bone 98 (2017) 59–67.

[37] M. Sinnesael, M.R. Laurent, F. Jardi, V. Dubois, L. Deboel, P. Delisser, G.J. Behets, P.C. D’Haese, G. Carmeliet, F. Claessens, D. Vanderschueren, Androgens inhibit the osteogenic response to mechanical loading in adult male mice, Endocrinology 156(4) (2015) 1343–53.

[38] T. Sugiyama, G.L. Galea, L.E. Lanyon, J.S. Price, Mechanical loading-related bone gain is enhanced by tamoxifen but unaffected by fulvestrant in female mice, Endocrinology 151(12) (2010) 5582–90.

[39] T. Sugiyama, L.K. Saxon, G. Zaman, A. Moustafa, A. Sunters, J.S. Price, L.E. Lanyon, Mechanical loading enhances the anabolic effects of intermittent parathyroid hormone (1-34) on trabecular and cortical bone in mice, Bone 43(2) (2008) 238–48.

[40] J.N. Farr, M. Xu, M.M. Weivoda, D.G. Monroe, D.G. Fraser, J.L. Onken, B.A. Negley, J.G. Sfeir, M.B. Ogrodnik, C.M. Hachfeld, N.K. LeBrasseur, M.T. Drake, R.J. Pignolo, T. Pirtskhalava, T. Tchkonia, M.J. Oursler, J.L. Kirkland, S. Khosla, Targeting cellular senescence prevents age-related bone loss in mice, Nat Med 23(9) (2017) 1072–1079.

[41] N. Holguin, M.D. Brodt, M.J. Silva, Activation of Wnt Signaling by Mechanical Loading Is Impaired in the Bone of Old Mice, J Bone Miner Res 31(12) (2016) 2215–2226.

[42] G.L. Galea, L.B. Meakin, M.A. Harris, P.J. Delisser, L.E. Lanyon, S.E. Harris, J.S. Price, Old age and the associated impairment of bones’ adaptation to loading are associated with transcriptomic changes in cellular metabolism, cell-matrix interactions and the cell cycle, Gene 599 (2017) 36–52.

[43] G. Zaman, L.K. Saxon, A. Sunters, H. Hilton, P. Underhill, D. Williams, J.S. Price, L.E. Lanyon, Loading-related regulation of gene expression in bone in the contexts of estrogen deficiency, lack of estrogen receptor alpha and disuse, Bone 46(3) (2010) 628–42.

[44] H.Z. Ke, T.A. Brown, H. Qi, D.T. Crawford, H.A. Simmons, D.N. Petersen, M.R. Allen, J.D. McNeish, D. D. Thompson, The role of estrogen receptor-beta, in the early age-related bone gain and later age-related bone loss in female mice, J Musculoskelet Neuronal Interact 2(5) (2002) 479–88.

[45] S.H. Windahl, O. Vidal, G. Andersson, J.A. Gustafsson, C. Ohlsson, Increased cortical bone mineral content but unchanged trabecular bone mineral density in female ERbeta(-/-) mice, J Clin Invest 104(7) (1999) 895–901.

